# Novel method for quantifying AhR-ligand binding affinities using Microscale Thermophoresis

**DOI:** 10.1101/2021.01.26.428246

**Authors:** Anne Stinn, Jens Furkert, Stefan H.E. Kaufmann, Pedro Moura-Alves, Michael Kolbe

## Abstract

The aryl hydrocarbon receptor (AhR) is a highly conserved cellular sensor of a variety of environmental pollutants and dietary-, cell- and microbiota-derived metabolites with important roles in fundamental biological processes. Deregulation of the AhR pathway is implicated in several diseases, including autoimmune diseases and cancer, rendering AhR a promising target for drug development and host-directed therapy. The pharmacological intervention of AhR processes requires detailed information about the ligand binding properties to allow specific targeting of a particular signalling process without affecting the remaining. Here, we present a novel microscale thermophoresis-based approach to monitoring the binding of purified recombinant human AhR to its natural ligands in a cell-free system. This approach facilitates a precise identification and characterization of unknown AhR ligands and represents a screening strategy for the discovery of potential selective AhR modulators.

## Introduction

Biomolecular interactions like protein-protein or protein-ligand interactions play an important role in almost all physiological processes. Remarkably, receptor agonist or antagonist interactions are of highest interest for the pharmaceutical drug development, where a detailed investigation of such processes is essential not only for understanding the underlying molecular mechanisms, but also the mode of action of a drug^1^. Quantitative binding studies can be performed by a number of biophysical approaches, such as surface plasmon resonance (SPR) or isothermal titration calorimetry (ITC)^2,3^. However, these techniques are often limited either due to the immobilization of one of the interaction partners that may interfere with binding (SPR) or due to high sample consumptions (ITC), respectively. Microscale thermophoresis (MST) is a relatively new method that allows for a fast and robust evaluation of biomolecular interactions in any desired buffer, including cell lysates or blood serum, without the need of surface immobilization or the use of excessive protein concentrations^4,5^.

The evolutionary highly conserved aryl hydrocarbon receptor (AhR) is a ligand-dependent transcription factor that mediates responses to environmental pollutants as well as dietary-, cell- and microbiota-derived metabolites^6^. Although originally identified as dioxin receptor, extensive research in the last years has demonstrated that the receptor is a key regulator of a broad spectrum of physiologically relevant functions, spanning from xenobiotic metabolism, developmental biology, as well as immunity^7–10^. Notably, its high relevance in the immune response of vertebrates, as well as the involvement in the onset of pathological conditions, e.g. cancer and autoimmune diseases, makes the AhR a promising target for drug development and host-directed therapy (HDT)^7,9,11^. As a member of the basic helix-loop-helix PER-ARNT-SIM (bHLH-PAS) protein family of transcriptional regulators, the AhR is characterized by a modular organization of regulatory domains, comprising an amino-terminal bHLH DNA binding domain that is contiguous to a tandem PAS A and PAS B domain^6,7^ (Fig 1a). The PAS B domain encompasses the receptors ligand-binding capability^6^. While the AhR domain architecture and the multifarious interactions with various proteins that enable the activation of different signalling pathways are well described^6,7^, little is known about the molecular processes of AhR ligand recognition. This is mainly because the crystal structure of the PAS B ligand-binding domain still remains to be solved. Moreover, although numerous AhR ligand binding analyses have been reported in the literature, in their majority radioligand competition assays and others are focusing on murine and rat AhR or species other than human^13,14,15^. Up to date, the lack of a robust recombinant expression system that allows for high-level production of functional human AhR has limited the development of a reliable and sensitive ligand binding assay that enables quantitative measurements of ligand binding affinities. Currently used methods (e.g. virtual ligand screening combined with AhR reporter cell studies or enzymatic activities) often result in divergent values, are time-consuming and do not allow for determination of direct binding, which is a critical aspect in drug discovery^16,17^. Therefore, we aimed to establish a robust protocol that allows for testing binding affinities of a purified recombinant human AhR to small molecules by using MST.

**Figure 1.**
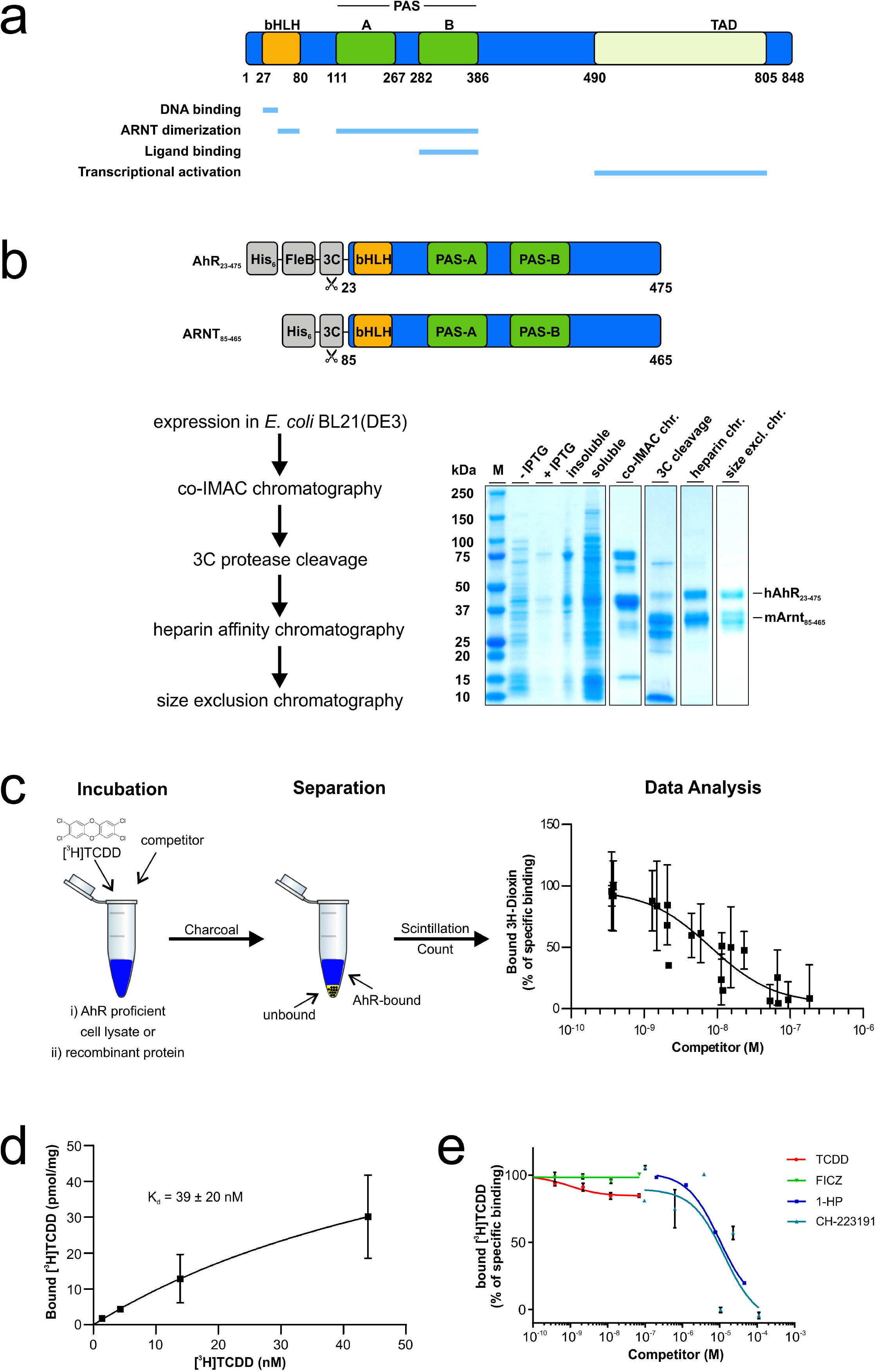
Verification of the ligand-binding properties of the recombinant AhR-ARNT by radioligand binding assay. (**a**) Functional domains of the human hAhR. Numbers indicate amino acid residues displaying the relative domain boundaries. (**b**) Schematic representation of the AhR-ARNT protein complex purification. Both proteins were co-expressed in *E. coli* BL21(DE3) and purified in a 3-step purification process. Samples from different steps of the purification were analyzed by SDS-PAGE and Coomassie staining. The theoretical molecular weight of the AhR and ARNT after cleavage with 3C is 51 and 40 kDa, respectively. (**c**) Principle of the radioligand binding assay. AhR-deficient and -proficient liver lysates were incubated with radioactively labelled [^3^H]TCDD and, in case of competition assays, in presence of increasing concentrations of unlabeled competitor ligands. After incubation, charcoal was added to remove unbound tracer molecules and radioactivity of the supernatant was measured. Specific binding was defined as the difference of radioactivity between extracts supplemented with recombinant AhR-ARNT and AhR-deficient lysates. (**d**) Saturation binding analysis of the purified AhR-ARNT receptor complex incubated with different concentrations of radioactively labelled [^3^H]TCDD. (**e**) Concentration-dependent displacement of [^3^H]TCDD from AhR-ARNT co-incubated with increasing concentrations of unlabeled competitor ligands. After 24 h, radioactivity in the supernatants was measured and competitive binding after NSB-subtraction was calculated. Data are mean ± SD of two independent experiments performed with triplicates.

## Results and discussion

### Recombinant AhR-ARNT is functionally active

In the past, numerous attempts to produce soluble AhR protein comprising the PAS B ligand-binding domain in sufficient amounts to support structural and functional studies failed^18,19^. Similarly, we observed that the expression of human AhR (hAhR) in *E. coli* and mammalian cells resulted exclusively in aggregated and/or insoluble protein material. This was irrespective of the expression construct and the AhR amino acid boundaries tested (data not shown). In contrast, several protocols describe the successful expression and purification of the aryl hydrocarbon nuclear translocator (ARNT)^20,21^. ARNT, also belonging to the bHLH-PAS protein family, sharing its typical domain architecture, is the interacting partner of AhR that was shown to dimerize and stabilize the AhR conformation upon ligand binding and receptor activation^6,21^. We, therefore, decided to make use of this properties and co-expressed the human AhR (hAhR_23-475_) comprising the ligand-binding domain in complex with murine ARNT (mARNT_85-465_) in *E. coli* BL21(DE3) (for details see Methods, Fig. 1b and Supplementary Fig. 1). Both proteins were expressed as C-terminal truncations as the C-terminal transactivation domain is not required for ligand binding and is known to have a tendency to aggregate^6^. To enhance AhR solubility upon bacterial expression, the protein was fused to an N-terminal FleB expression tag derived from the *Yersinia enterocolitica* flagellin FliC/FljB, which was shown to be monomeric and expressed at high levels in *E. coli*^22^. Noteworthy, fusion to the N-terminal solubility tag together with expression in presence of mARNT not only enhanced the expression levels of hAhR but also resulted in a significantly increased solubility of the receptor protein (Fig. 1b). This ultimately allowed us to purify sufficient amounts of recombinant hAhR in complex with mARNT to be further subjected to functional analyses (Fig. 1b, Supplementary Fig. 1). We initially validated the functionality of the purified protein complex employing the conventional cell-based radioligand binding assay^18^ (Fig. 1c). In this, an equilibrium dissociation constant (K_d_) of 2.7-4.2 nM was ascertained for the bona fide ligand 2,3,7,8-tetrachlorodibenzodioxin (TCDD) to the endogenous hAhR in hepatocyte cell lysates^13^. We assessed the specific binding of radioactively labelled [^3^H]TCDD tracer to the recombinant AhR-ARNT receptor in the presence of liver lysates derived from AhR deficient mice. Saturation binding experiments verified the direct binding of the prototypical ligand TCDD to the purified AhR-ARNT complex with a calculated K_d_ of 39 ± 20 nM, and a maximal binding capacity (B_max_) of 60.3 ± 17.3 pmoles/mg protein (Fig. 1d). The observed binding was specific to hAhR, as we did not detect any interaction between the individually purified mARNT protein and [^3^H]TCDD (Supplementary Fig. 2)^20^.

Next, we performed competition radioligand binding experiments to evaluate the binding ability of the recombinant AhR-ARNT to diverse AhR ligands belonging to different structural classes (Fig. 1e). The bacterial pigment 1-hydroxyphenazine (1-HP) from Pseudomonas aeruginosa, previously identified by our group as a potent AhR activator^8^, was chosen as a representative of microbe-derived AhR ligands. CH-223191, a commercially available AhR-inhibitor^23^, was chosen as AhR antagonist. We observed a concentration-dependent displacement of the tracer for both 1-HP and CH-223191, with specific K_i_’s of 9.2 μM (95% confidence limit: 7.4-11.1 μM) and 12.2 μM (95% confidence limit: 1.8-76.1 μM), respectively, validating the functional activity of the recombinant AhR-ARNT complex in terms of its capability to bind small molecules. Further, we tested the binding affinity of the endogenous AhR agonist 6-formylindolo-[3,2-b]-carbazole (FICZ)^6,7,24^ and the prototypical exogenous ligand TCDD to AhR in competition binding assays (Fig. 1e). Though reported to have an affinity in the pM-nM range, we failed to see competition when incubating the purified AhR-ARNT with increasing concentrations of unlabeled FICZ, which was likely attributed to the high instability of its tryptophan group in aqueous solution^25^. However, in agreement with the data obtained from our radioligand saturation experiments, we observed only an initial competition of unlabeled TCDD for binding to hAhR at the highest concentration. Full displacement of the radioligand would require the use of even higher concentrations of unlabeled TCDD, which due to its insolubility in water^26^ is not feasible in this experimental set-up.

### MST is a sensitive method to determine AhR-ligand binding affinities

The radioligand binding assay, although commonly used, is known to be extremely sensitive to minor changes in the protocol that likely lead to marked differences in the measured affinities^13,18,26^. Here, total cytosolic extracts of hepatocytes are being used; therefore, the observed ligand binding may not be specific to AhR, but potentially also arise from unspecific interactions of the ligand with other constituents of the cytosol. Additionally, due to high background noise in this assay, only subtle differences in radioactivity can be measured^8,18^. Thus, the limit and detection range for analyzing specific binding is quite small, which urges the development of an improved AhR ligand binding assay that can replace the classical radioligand method.

We implemented an MST based assay, which in addition to the low protein amount required and short assay times, is also characterized by its buffer independency, enabling the analysis of the receptor under close-to-native conditions. Exploiting the intrinsic fluorescence of proteins, it allows the analysis of biomolecular interactions without the need for sample modifications due to labelling (Fig. 2a)^5,27^. To evaluate the feasibility of the label-free MST approach for analyzing small molecule binding to AhR, we first tested the binding of TCDD to purified AhR-ARNT complexes (Fig. 2b). We titrated the non-fluorescent ligand to a given concentration of AhR-ARNT complex (250 nM final). Concentration-dependent ligand-induced differences were readily recorded in the MST traces using AhR-ARNT, but not when mARNT was used alone, validating the specific binding of TCDD to hAhR (Supplementary Fig. 3). AhR binding affinity to TCDD was calculated (K_d_ of 139 ± 99 nM) (Fig. 2b). Next, we elucidated the compatibility of MST for analyzing binding of the recombinant AhR-ARNT to small molecules of various classes (Fig. 2c-e). In contrast to the radioligand binding assay, we were able to measure specific saturable binding of hAhR to the labile high-affinity agonist FICZ with a computed binding affinity of 79 ± 36 nM. Employing the same setup, we further confirmed binding of 1-HP and CH-223191 with binding affinities in the higher nM range with K_d_’s of 943 ± 176 nM and 496 ± 82 nM, respectively. Together, this validated the suitability of the devised approach for measuring binding of recombinant AhR-ARNT complexes not only to AhR activators such as 1-HP (Fig. 2c,d) but also to specific inhibitors such as the CH-223191 (Fig. 2e).

**Figure 2.**
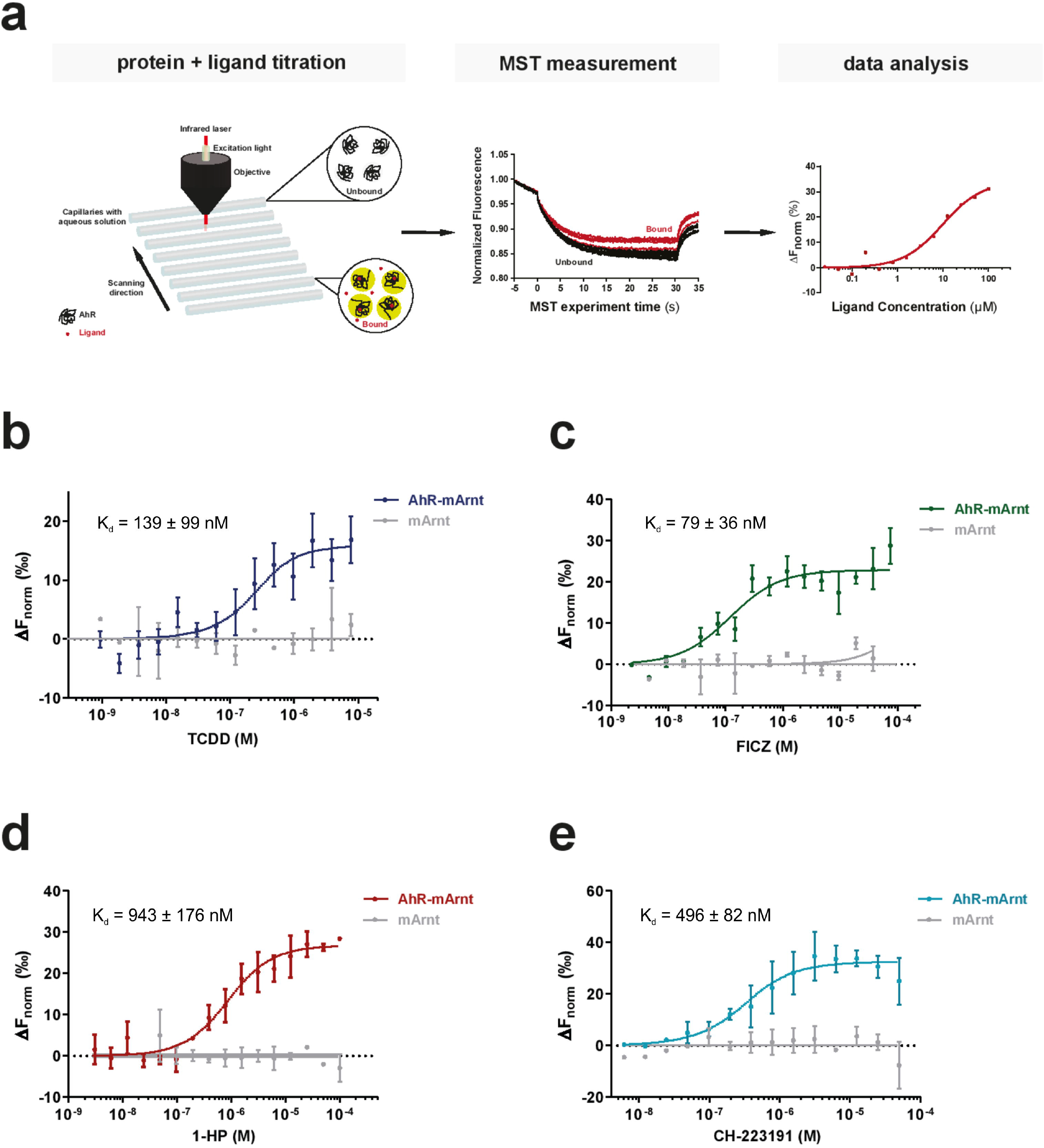
Suitability of MST as a sensitive method to determine AhR-ligand binding affinities. (**a**) The basic principle of an MST ligand binding experiment. The thermophoretic movement of the intrinsically fluorescent protein in the presence of increasing concentrations of a non-fluorescent ligand is analysed. In the case of binding (bound, red traces), thermophoresis will differ from the unbound state and will result in a gradual change in the recorded MST traces. Plotting the changes in the normalized fluorescence as a function of the ligand concentration will yield a binding curve that can be fitted to calculate the binding affinities. (**b-e**) Binding of the prototypic ligand TCDD (**b**), the endogenous ligand FICZ (**c**), the bacterial pigment 1-HP (**d**) and the AhR inhibitor CH-223191 (**e**) to 250 nM recombinant AhR-ARNT complex was analysed with label-free MST (coloured lines). To verify that the measured interaction was exclusive for AhR, we examined binding to separately purified ARNT (grey line). Error bars represent the ± SD of each tested ligand concentration calculated from three independent experiments.

## Conclusion

Traditionally, AhR binding assays have been done using radiolabeled ligands as the method of choice^13,14,18^. However, not only because of the undesirable usage of radioactive and toxic tracer material, the use of animals for organ extraction and the increased costs involved but also due to diverse technical reasons, there is a necessity for the development of an improved ligand binding assay. Besides the apparent sensitivity towards amendments in the protocol, some studies stress the fact that the radioligand binding assay largely suffers from the insolubility of TCDD and other AhR ligands in water^25,26^. Due to their lipophilic nature, these compounds are usually dissolved in DMSO. At high concentrations, dilution with the assay buffer results in precipitation and unspecific interactions of the small molecule candidates with hydrophobic surfaces, thereby affecting the final results. Yet, the development of a precise and sensitive AhR ligand binding assay was limited due to the lack of an expression system yielding high amounts of functional AhR protein. This study demonstrates that the recombinant expression of hAhR fused to an N-terminal FleB-tag and in complex with its interacting partner ARNT ultimately allows the purification of significant amounts of functionally active human receptor protein with yields of 1 mg protein per litre expression culture. Further, the purified protein is suitable for analyzing receptor-ligand interactions using MST, which provides a powerful alternative to the classical radioligand binding assay. Exploiting the protein’s intrinsic fluorescence free in solution obviates the need for sample modifications like labelling or immobilization that might interfere or affect ligand binding. Using this approach, we confirmed the binding of hAhR to a known, *bona fide* ligands, including TCDD, FICZ, 1-HP and CH223191. Yet, compounds with an intrinsic fluorescence at or close to λ_ex_ ~ 280 nm cannot be tested with this method due to the overlapping fluorescence with the target protein. The direct fusion of the target protein to a fluorescent tag, e.g. GFP or RFP that can be specifically monitored at the Monolith NT.115 machine, would circumvent this limitation, make additional purification steps needless and thus allow measurements to be done directly in cell lysates^28^.

Though the obtained binding affinities for the individually tested ligands to the recombinant protein are slightly lower than the ones reported in the literature for the hAhR^13,24,26^, such differences are presumably explainable by missing endogenous factors that are likely to support ligand sensing of the cytosolic AhR, e.g. stabilizing effects of chaperones (HSP90, XAP2) in the unbound receptor state^29,30^. Moreover and of note, most of the literature-described binding affinities were experimentally determined for the murine AhR, which is known to display an almost 10-fold higher binding affinity towards TCDD than human AhR^31^.

Finally, the considerable advantages of buffer independency and the short reaction times enable fast and reliable monitoring of AhR binding to a large number of small molecules, including exogenous AhR ligands, AhR inhibitors, and also to less stable compounds (e.g. FICZ), for which previous interaction studies with longer assay times have proven to be challenging^18,24^. Altogether, in addition to the implementation of this method in basic research for analyzing AhR-ligand interactions and the signalling pathway in more detail, the novel AhR binding assay presented here bears the potential for future drug/compound screenings. Ultimately, this will facilitate the discovery of potential selective modulators of this intriguing broad-spectrum receptor of high interest for HDT.

## Material and Methods

### Resource availability/contact for Reagent and Resource sharing

Further information and requests for resources and reagents should be directed to and will be fulfilled by the corresponding author, Pedro Moura-Alves or by the co-correspondent author, Michael Kolbe.

### Ligands

All compounds were obtained from commercial sources, solubilized in dimethyl sulfoxide (DMSO) and stored protected from light. 2,3,7,8-Tetrachlorodibenzo-p-dioxin (TCDD) was obtained from LGC Standard. [^3^H]TCDD diluted in ethanol was from Hartmann Analytic. 1-hydroxyphenazine (1-HP) was purchased from TCI Europe N.V., 6-Formylindolo[3,2-b]carbazole (FICZ) and 2-Methyl-2H-pyrazole-3-carboxylic acid (2-methyl-4-o-tolylazophenyl)-amide (CH-223191) from Sigma-Aldrich. 1-HP, CH-223191 and FICZ were kept at RT, 4°C or −20°C, respectively.

### Plasmid construction and protein preparation

For the recombinant expression in *E. coli*, a codon-optimized fragment of human *AhR* (UniProt ID: P35869) encompassing the bHLH-PAS A-PAS B domains (amino acid residues 23 to 475) was commercially synthesized (MWG Eurofins) and cloned into pET21b harbouring an N-terminal His_6_- and the *Yersinia*-derived flagellin subunit FleB (amino acid residues 54 to 332, UniProt ID: A1JSQ5) as expression tag, followed by an HRV 3C protease cleavage site. The resulting construct was confirmed by DNA sequencing. The pET30-EK/LIC mARNT expression plasmid encoding the murine ARNT residues 85 to 465 (Δ274-297, C256S, Δ351-358, UniProt ID: P53762) was a generous gift from Prof. Dr. Oliver Daumke (MDC Berlin). The amino acid sequences of the expression vectors are shown in Supplementary Fig. 4. Both plasmids were co-transformed into *E. coli* BL21(DE3) competent cells (Novagen). Bacteria were grown to an OD_600_ of 0.8 in lysogenic broth (LB (Luria/Miller)) medium (Carl Roth) before the expression was induced with 0.5 mM isopropyl-β-D-thiogalactopyranoside (IPTG, Sigma-Aldrich). Following overnight expression at 18°C, proteins were purified as described previously^12,21^ (see also Supplementary Fig. 1). Shortly, pellets were lysed using an SLM Aminco Pressure Cell Press and the clarified lysate was applied onto a HisTALON Superflow column (Clontech). Bound proteins were eluted by applying an increasing imidazole concentration, buffer exchanged and N-terminal His_6_-tags were removed with HRV 3C protease (3C) overnight at 4°C. The cleaved protein complex was further purified on a HiTrap Heparin HP column (GE Healthcare), followed by size exclusion chromatography on a Superdex 200 10/300 GL equilibrated in 20 mM HEPES pH 8.0, 200 mM NaCl, 5% glycerol, and 5 mM DTT. Peak fractions containing AhR-mARNT were pooled and concentrated using Amicon filter units (Millipore).

Mouse ARNT (residues 85-465) was expressed as N-terminal His_6_-fusion protein in *E. coli* BL21(DE3) and purified as described above for the heterodimer complex.

### Radioactive labelled TCDD competition

Radioligand binding assays were performed as previously established^13^. Briefly, for saturation experiments, 0.5 mg/ml of purified AhR-ARNT or ARNT protein was diluted in MDEG buffer (25 mM MOPS, 1 mM DTT, 1 mM EDTA, 10% glycerol, 20 mM molybdate, pH 7.5) and incubated with increasing concentrations of radioactively labelled [^3^H]TCDD tracer. Reactions were supplemented with liver lysate of AhR-deficient mice (AhR^−/−^, C57BL/6 background) to a final protein concentration of 5 mg/ml to reduce unspecific binding and adsorption of the tracer to plastic surfaces. Liver lysates were prepared in MDEG buffer by homogenization using the gentleMacs Dissociator (Miltenyi Biotec), followed by ultracentrifugation at 100,000xg and 4°C for 1h. Cytosolic fractions were collected, protein concentration determined by Bradford reaction (Protein Assay Kit, Pierce), and further diluted to a final protein concentration of 5 mg/ml in MDEG buffer. Binding reactions were incubated at 4°C for 24 h in order to ensure equilibrium conditions. Subsequently, 30 μl of a Norit A charcoal suspension (100 mg/ml equilibrated in MDEG buffer) was added into each 200 μl of the reaction mixture. Samples were kept on ice for 15 min and centrifuged for another 15 min at 25,000xg and 4°C. Following this, 150 μl of the supernatant was carefully transferred and radioactivity was measured in a liquid scintillation counter (Tri-Carb 3110TR, PerkinElmer). All samples were measured in triplicate. For competition assays, a constant concentration of [^3^H]TCDD (4 nM final) was used, to which a serial dilution of unlabeled competitor molecule dissolved in DMSO was added in excess. All subsequent steps were carried out as described above. The specific binding was defined as the difference of radioactivity between the reactions supplemented with the recombinant protein and AhR-deficient (Ahr^−/−^) mouse lysates. Analysis of the data and calculation of the binding constants K_d_ and B_max_ was performed with GraphPad Prism 7 by using a nonlinear regression fitting a saturation binding isotherm. For competition studies, the IC50 values were obtained by fitting a one-site competitive binding equation to the experimental data. K_i_ values were derived from IC50 using the Cheng-Prusoff equation^32^.

### Microscale Thermophoresis (MST)

Microscale thermophoresis (MST) experiments were performed according to the manufacturer’s instructions (NanoTemper Technologies GmbH) using a Monolith NT.LabelFree. In brief, 250 nM of purified AhR-ARNT proteins were diluted in assay buffer (20 mM HEPES pH 8.0, 200 mM NaCl, 5% glycerol, 5 mM DTT, and 0.1% Pluronic F-127) and incubated for 5 min, in the presence of a serial dilution of the different ligands (ligand solubilized in DMSO at a final constant concentration of 2%). After incubation, samples were filled into NT.LabelFree Zero Background MST Premium coated capillaries (NanoTemper Technologies GmbH) and measurements were taken, at a constant temperature of 22°C. MST traces were collected with an LED excitation power of 20% and an MST laser power of 20% and 40%. The MO.Control Analysis software (NanoTemper) was used to analyze the interaction affinity and the dissociation constant (K_d_) for each ligand using the Kd fit model. Changes in the normalized fluorescence (ΔFnorm [‰]) were plotted as a function of the ligand concentration.

### Mice handling for preparation of liver cell suspensions

AhR-deficient mice (*Ahr*^−/−^, C57BL/6 background) were kindly provided by B. Stockinger (The Francis Crick Institute, London, United Kingdom). *Ahr*^−/−^ mice were bred in the Max Planck Institute for Infection Biology mouse facility. Animal experiments were carried out according to institutional guidelines approved by the local ethics committee of the German authorities (Landesamt für Gesundheit und Soziales Berlin; Landesamt für Verbraucherschutz und Lebensmittelsicherheit, project number G0257/12). Liver were isolated from mice of 8– 12 weeks of age following published protocols^8^.

### Quantification and statistical analysis

Data fitting including baseline corrections, normalization, calculation of mean and error (SEM), and statistical tests were carried out in GraphPad Prism (version 7). The number of biological and technical replicates, as well as the entity plotted, are indicated in the figure legends. Data baseline correction and normalization, where applied, were indicated in the corresponding method section and the axis labels. No explicit power analysis was used.

## Data Availability

The authors declare that this study did not generate or analyze a datasets/code.

## Acknowledgements

The authors acknowledge those who have provided tools and materials for this work. Brigitta Stockinger (The Francis Crick Institute, London, UK) for the *Ahr*^−/−^ mice. Expression plasmid pET30-EK/LIC-ARNT was a gift from Oliver Daumke and Kathrin Schulte (Max Delbruck Center, Berlin, Germany). We acknowledge the lab of Rainer Haag (Freie Universität Berlin, Berlin, Germany) for providing access to Monolith NT.LabelFree, Maria Garcia Alai, Ioana Maria Nemtanu and Janina Hinrichs (European Molecular Biology Laboratory, Hamburg, Germany) for technical assistance and Juana de Diego (Center for Structural Systems Biology, Hamburg, Germany) for scientific discussions.

This work was generously supported by intramural funding of the Max Planck Society to SHEK, by the International Max Planck Research School for Infectious Diseases and Immunology (IMPRS-IDI) and by iNEXT, infrastructure for NMR, EM and X-rays for Translational Research, project number 653706, funded by the Horizon 2020 programme of the European Union.

## Author contributions

AS, SHEK, PM-A and MK conceived and AS and MK designed the study. AS designed and performed the experiments and data analysis. AS and JF performed the radioligand binding studies. AS, PM-A, MK wrote the manuscript. All authors commented on the paper.

