## Supplementary Information for "Novel method for quantifying AhR-ligand binding affinities using Microscale Thermophoresis"

**Affiliations:**
[1]Department of Immunology, Max Planck Institute for Infection Biology, Charitéplatz 1, 10117 Berlin, Germany. [2]Department of Structural Infection Biology, Center for Structural Systems Biology (CSSB), Helmholtz-Center for Infection Research (HZI), Notkestrasse 85, 22607 Hamburg, Germany. [3]Leibniz-Institut für Molekulare Pharmakologie (FMP), Robert-Rössle-Strasse 10, 13125 Berlin, Germany. [4]Hagler Institute for Advanced Study at Texas A&M University, College Station, TX 7843. [5]Max Planck Institute for Biophysical Chemistry, Am Faßberg 11, 37077 Göttingen, Germany. [6]Ludwig Institute for Cancer Research, Nuffield Department of Clinical Medicine, University of Oxford, OX3 7DQ Oxford, UK. [7]Faculty of Mathematics, Informatics and Natural Sciences, University of Hamburg, Rothenbaumchaussee 19, 20148 Hamburg, Germany.

**Supplemental information:**

**Fig. S1 - Purification of recombinantly expressed AhR-ARNT protein complex.**

**Fig. S2 - Radioligand assay of AhR-ARNT and ARNT.**

**Fig. S3 - Raw data of MST experiments for AhR-ARNT and ARNT.**

**Fig. S4 - Expression vectors used in this study and the encoded protein amino acid sequences.**

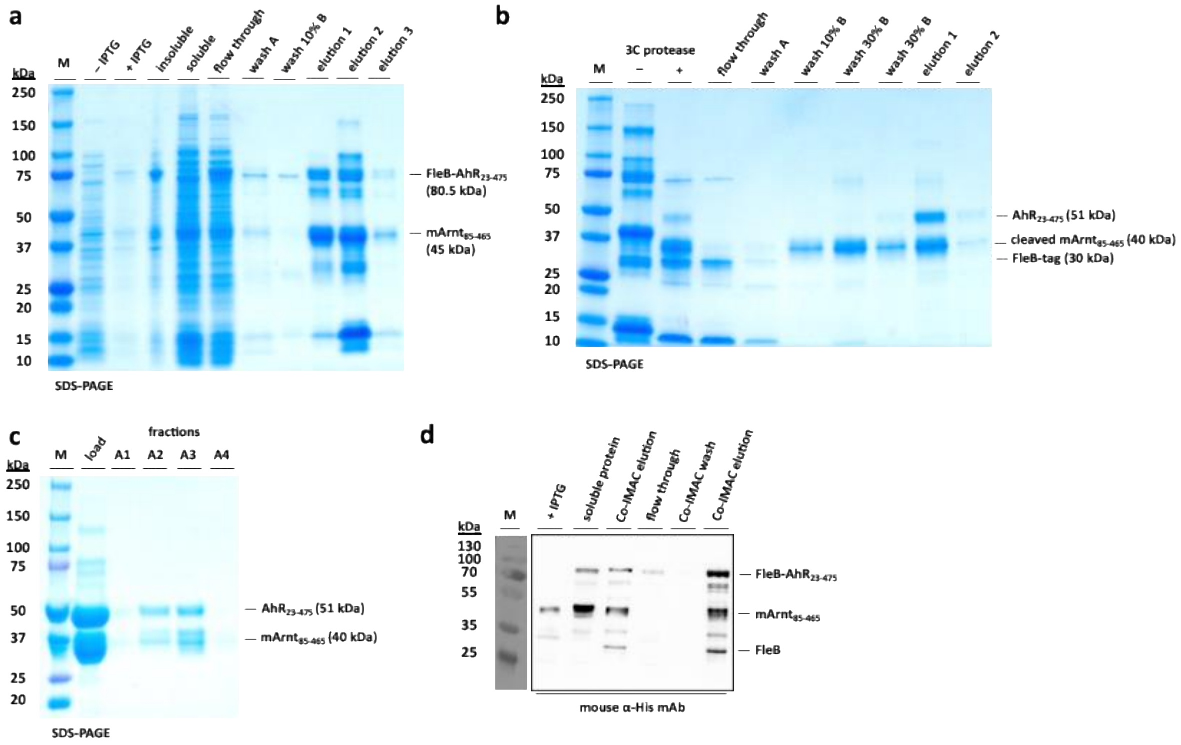


**Fig S1: Purification of recombinantly expressed hAhR-mARNT protein complex.**

AhR-ARNT protein complexes were generated by co-expression in *E. coli* BL21(DE3) and subsequent purification of soluble material in 3 steps. Protein fractions collected throughout the purification process were analyzed by SDS-PAGE and Coomassie-staining. (**a**) His-tagged proteins were pulled down from the bacterial lysate via affinity chromatography using Cobalt-IMAC resin. Elution fractions containing FleB-hAhR$_{23\text{-}475}$ and mARNT$_{85\text{-}465}$ were pooled and incubated with HRV3C protease. (**b**) Following tag cleavage, the AhR-ARNT protein complexes were further purified via heparin affinity chromatography. (**c**) For final polishing, the AhR-ARNT complexes were separated from aggregates on a Superdex S200 column. (**d**) Western blot analysis of the purified FleB-hAhR$_{23\text{-}475}$ and mArnt$_{85\text{-}465}$ using an anti-His-tag monoclonal antibody.

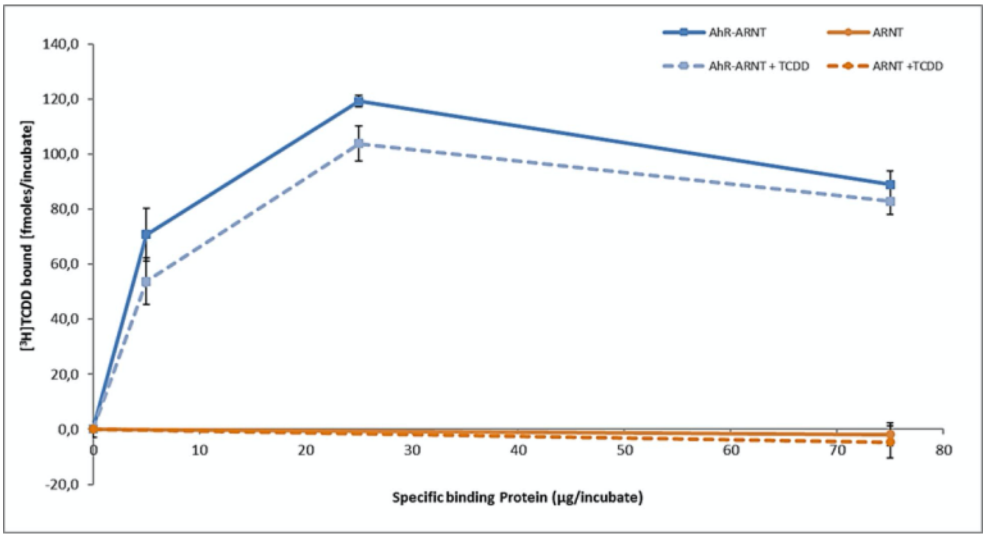


**Fig S2: Radioligand assay of hAhR-mARNT and mARNT.**

The AhR-ARNT complex and the separately purified mARNT protein were tested for their ligand binding properties by performing the radioligand binding assay. For this, increasing concentrations of protein (in µg) were incubated for 48 h with constant concentrations of radioactively labeled [$^{3}$H]TCDD in the presence or absence of unlabeled competitor (TCDD) prior to the measurement of radioactivity. Specific binding was calculated after the subtraction of non-specific binding (NSB) derived from AhR knockout (Ahr$^{-/-}$) liver lysates. While there was a clear binding of [$^{3}$H]TCDD and displacement by unlabeled dioxin detectable for the AhR-ARNT complex, no binding was observed for ARNT. Protein concentration (µg/reaction) on X-axis and bound fraction on the Y-axis are shown. Results are derived from multiples experiments repeated in triplicates.

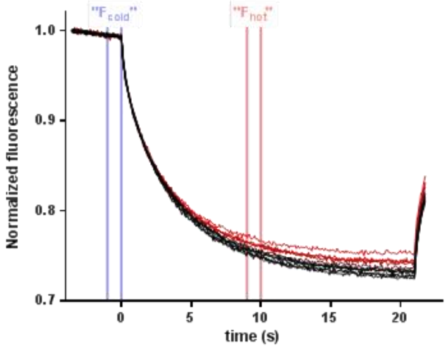


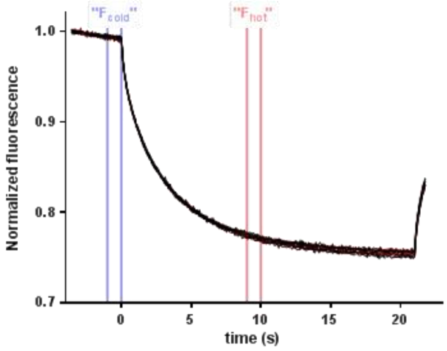


**Fig S3: Raw data of MST experiments for hAhR-mARNT and mARNT.**

In the MST experiment, serial dilutions of ligand dissolved were added to constant protein concentrations of recombinant AhR-ARNT or mARNT. For analysis, MST traces for each protein-ligand reaction were collected where the changes in the normalized fluorescence were plotted as a function of time. While for AhR-ARNT concentration-dependent ligand-induced differences were recorded, the MST traces for mARNT did not show concentration-dependent variations, indicative for no protein-ligand interaction.

**pET28a-His$_6$-FleB-<u>3C</u>-hAhR$_{23-475}$**

MGSSHHHHHHSSGLVPRGSHMTSNINGLTVAARNANDGISLSQTAEGALGEINNNLQRVRDL
TVQAQNSSNSASDIDSIQSEVNQRMEEINRVTKQTDFNGIKVLDNRTKTDSSYDFQVGSKDN
EQISIAIGASSGWNLATANADGTSSDTVNTYAFTKKAALDTAQTDYDTANTAYLAAVKSGVA
GDITTTKATLDGKNTALATAVKDATAVNEAVNGKVRTVAAKGFDVLNGTVAADGKATGTTPL
ADIDKALKAVDTQRSVLGASQNRFESTITNLNNTVNNLTSARGGASGGGS<u>LEVLFQGP</u>MVKP
IPAEGIKSNPSKRHRDRLNTELDRLASLLPFPQDVINKLDKLSVLRLSVSYLRAKSFFDVAL
KSSPTERNGGQDNCRAANFREGLNLQEGEFLLQALNGFVLVVTTDALVFYASSTIQDYLGFQ
QSDVIHQSVYELIHTEDRAEFQRQLHWALNPSQCTESGQGIEEATGLPQTVVCYNPDQIPPE
NSPLMERCFICRLRCLLDNSSGFLAMNFQGKLKYLHGQKKKGKDGSILPPQLALFAIATPLQ
PPSILEIRTKNFIFRTKHKLDFTPIGCDAKGRIVLGYTEAELCTRGSGYQFIHAADMLYCAE
SHIRMIKTGESGMIVFRLLTKNNRWTWVQSNARLLYKNGRPDYIIVTQRPLTDEEGTEHLRK
RNTKLPFMFTTGEAVLYEATNPFPAIMDPLPLRTKNGTSGKDSATTSTLSKDSLNPSSLLAA
MMQQDESIYLYPASST

**pET30 EK LIC-His$_6$-<u>3C</u>-mARNT$_{85-465\ \Delta 274-297\ C256S\ \Delta 351-358}$**

MHHHHHHSSGLVPRGSGMKETAAAKFERQHMDSPDLGTDDDDKM<u>LEVLFQGP</u>GSDPDKERLA
RENHSEIERRRRNKMTAYITELSDMVPTCSALARKPDKLTILRMAVSHMKSLRGTGNTSTDG
SYKPSFLTDQELKHLILEAADGFLFIVSCETGRVVYVSDSVTPVLNQPQSEWFGSTLYDQVH
PDDVDKLREQLSTSENALTGRVLDLKTGTVKKEGQQSSMRMSMGSRRSFICRMRCGTSSEGE
PHFVVVHCTGYIKAWPPAGVSLPDDDPEAGQGSKFCLVAIGRLQVTSSPNQPTEFISRHNIE
GIFTFVDHRCVATVGYQPQELLGKNIVEFCHPEDQQLLRDSFQQVVKLKGQVLSVMFRFRSK
TREWLWMRTSSFTFQNPYSDEIEYIICTNTNVK

**Fig S4. Expression vectors used in this study and the encoded protein amino acid sequences.**
